# Multi-domain Brain Age from Digital Cognitive Metrics as a novel approach for new longevity

**DOI:** 10.64898/2026.04.30.721651

**Authors:** Marta Arbizu Gómez, Carolina Sastre-Barrios, Elina Maltseva, Jorge M. Corada, César Ortea Suárez, íñigo Fernández de Piérola, Genny Lubrini, José A. Periáñez, Marcos Ríos-Lago, Jesús M. Cortés

## Abstract

**Background:** The continuous rise in life expectancy introduces a central challenge of new longevity, ensuring that the additional years gained are accompanied by the preservation of cognitive function and quality.

**Methods:** We propose a modeling framework for multi-domain brain age derived from a repertoire of digital cognitive metrics. The model, based on Ridge regression with Leave-One-Out cross-validation, was trained in a cohort of 394 healthy controls (HC; 307 women and 87 men; mean age 30.0 ± 12.5 years; range 17–64).

**Results:** The model achieved a correlation between chronological age and predicted age of r = 0.942 with a mean absolute error of 3.05 years. When applied to three additional clinical cohorts, multiple sclerosis (N = 70), traumatic brain injury (N = 23), and depression (N = 18), the model detected significant accelerated cognitive aging across all conditions, with processing speed emerging as the dominant contributor to accelerated aging, albeit with varying degrees of concentration across pathologies.

**Conclusions:** Digital cognitive metrics provide an accessible, non-invasive, and scalable biomarker for tracking brain aging, with strong potential for informing personalized neuropsychological interventions and for integration into active aging frameworks within the context of modern longevity.

## Introduction

The so-called *new longevity* is no longer a demographic projection, but a reality that is transforming the social, economic, and healthcare structure of contemporary societies. Spain, with a life expectancy at birth exceeding 83 years, ranks among the longest-lived countries in the world [1]. However, the extension of lifespan does not in itself guarantee that these additional years will be lived under conditions of well-being. In particular, the preservation of cognitive health emerges as one of the central challenges of this new stage of life: the prevalence of mild cognitive impairment and dementia increases exponentially with age, and the number of people affected by dementia in Europe is estimated to double over the next three decades [2].

Against this background, it is essential to have tools capable of monitoring individuals’ cognitive trajectories throughout the lifespan, detecting early deviations from normative aging, and implementing preventive interventions before decline becomes clinically manifest [3]. In this sense, the concept of life expectancy is incomplete unless it is complemented by the notion of *healthy life expectancy*, which refers to a life free from significant disease or disability [4]. This concept highlights the need for early detection of subtle cognitive decline in order to initiate compensatory intervention plans that prolong well-being and quality of life for as long as possible [5].

Over the past decade, the concept of *brain age* has become established as a reference framework for quantifying the discrepancy between the biological age of the brain and an individual’s chronological age [3]. It was originally developed in the field of neuroimaging through machine learning models trained on magnetic resonance imaging data from healthy individuals, allowing estimation of the age “imprinted” in a person’s brain based on its structure or connectivity [6]-[8]. The difference between estimated brain age and chronological age, commonly referred to as the brain age delta (Δ) or gap, functions as a biomarker. Thus, a positive delta indicates accelerated aging, with a brain age greater than chronological age, whereas a negative delta or one close to zero suggests a preserved or even “rejuvenated” aging trajectory.

The neuroimaging-based brain age paradigm has generated a robust body of evidence. The brain age gap (Δ) has been associated with mortality, progression of neurodegenerative diseases, treatment response, and cardiovascular risk factors, among others [9]-[11], and has demonstrated clinical utility in numerous neurological and psychiatric conditions [12]-[15]. However, neuroimaging-based models present important limitations for large-scale implementation: they require expensive equipment, specialized technical personnel, and hospital infrastructure, as well as harmonization procedures across scanners and even imaging sequences, all of which restrict their use to clinical and research contexts. As a consequence, neuroimaging-based brain age models have shown limited scalability, with MRI scans acquired on different machines often yielding different brain age estimates.

More recently, several authors have explored the possibility of estimating brain age from behavioral and cognitive data [16]. In particular, AgeML [17], an open-source framework for age modeling with machine learning, enables application of the brain age paradigm to any type of quantitative variable, including cognitive metrics. This approach opens the door to more accessible and cost-effective brain age models without sacrificing the ability to detect individual deviations from normative aging.

Digital neuropsychological assessment offers a complementary avenue to address this limitation [18], [19]. Computerized cognitive batteries provide objective, standardized, and high temporal resolution measurements (on the order of milliseconds) across multiple cognitive domains, at a significantly lower cost and with much greater scalability than neuroimaging. These characteristics make them ideal candidates for supporting cognitively based brain age models that can be deployed in a wide range of settings: primary care consultations, public health programs, telemedicine platforms, or even digital health applications.

In this study, we present a multicognitive brain age model trained exclusively on 18 digital cognitive metrics from the SPAIn battery (Speed of Processing Assessment Instrument [20]). Our objectives are threefold: **(a)** to demonstrate that digital cognitive metrics accurately capture the normative trajectory of cognitive aging; **(b)** to apply the model to three clinical cohorts with known accelerated cognitive aging—multiple sclerosis, traumatic brain injury, and depression—to assess its sensitivity; and **(c)** to discuss the potential of this approach as a tool for monitoring cognitive aging in the context of the new longevity and for designing personalized interventions.

## Methodology

### Participants

The study included four independent cohorts. The training cohort consisted of 394 healthy controls (HC; 307 women and 87 men; mean age 30.0 ± 12.5 years; range 17–64), defined as individuals with no known diagnosis of neurological disease, psychiatric disorder, or cognitive impairment at the time of assessment. The predominant educational level was university education (68.5%), followed by secondary education (9.9%), postgraduate studies (8.4%), doctoral studies (8.4%), and primary education (4.8%). The three clinical cohorts included patients with relapsing-remitting multiple sclerosis (MS; N = 70; 47 women and 23 men; mean age 41.2 ± 8.0 years; range 26–60), traumatic brain injury (TBI; N = 23; 17 women and 6 men; mean age 32.7 ± 12.7 years; range 17–64), and major depression (MD; N = 18; 13 women and 5 men; mean age 45.5 ± 8.1 years; range 31–58).

The Ethics Committee of the institution approved the study. Subjects were informed about the purpose of the investigation before the experimental session and signed a consent form according to the Declaration of Helsinki.

### Instrument: SPAIn Battery

All participants completed the SPAIn processing speed assessment battery [20], administered in digital format. This is a digital neuropsychological assessment instrument designed to measure multiple components of processing speed, attention, and executive functions. The relevance of these domains to the study of aging is well established. Processing speed is one of the cognitive functions that shows the earliest and most pronounced decline with age [21], and its deterioration has been associated with difficulties in daily life, loss of functional autonomy, and increased risk of subsequent cognitive decline [22]. Executive functions, in turn, constitute a central axis of cognitive aging and have proven sensitive to both normative and pathological aging.

For the present study, 18 quantitative metrics were extracted from four selected subtests, as described below. From the **Finger Tapping** subtest, four variables were obtained: mean number of taps in 10 seconds (left and right hand) and mean inter-tap interval (left and right hand), reflecting motor speed and lateralization. From the **Simple Reaction Time** subtest, three variables were derived: number of correct responses, number of omissions, and mean reaction time for correct responses, capturing elementary processing speed and basic attentional capacity. From the **Go/No-Go** subtest, four variables were extracted: correct responses in Go trials, false alarms in No-Go trials, omissions in Go trials, and mean reaction time for correct Go responses, assessing sustained attention and response inhibition. Finally, from the **Response Selection** subtest, seven variables were obtained: correct responses, commission errors, omissions, mean reaction time for correct responses, mean reaction time for commission errors, and post-error slowing, reflecting executive functioning, inhibitory control, and self-regulation capacity.

For the clinical cohorts, analyses were conducted using the subset of variables overlapping with those available in the healthy control cohort: 15 variables in MS, 5 in MD, and 18 in TBI.

### Brain Age Modeling

A regularized Ridge regression model was used to predict chronological age from the 18 cognitive metrics. Ridge regularization was selected because of its robustness to multicollinearity among predictors, a characteristic expected in cognitive metrics that share underlying processes, similar to the implementation described in [17]. The model was trained exclusively on the healthy control cohort using Leave-One-Out cross-validation (LOOCV), a procedure particularly suitable for addressing overfitting and evaluating the model’s generalization capacity in moderately sized samples by maximizing the use of available data.

Once brain age modeling was trained on healthy controls, the model was applied to the three clinical cohorts to obtain the estimated brain age of each patient. Brain age delta was calculated as the difference between estimated brain age and chronological age (Δ = estimated age − chronological age). A bias correction was applied following the procedure described in the AgeML framework [17], in order to correct the well-known tendency of brain age models to overestimate age in younger individuals and underestimate it in older individuals.

Mean delta was compared between healthy controls and each clinical cohort using a non-parametric two-sample (unpaired) test; p<0.05 was considered indicative of significant group differences. In addition, the relative contribution of each cognitive variable to delta was analyzed across pathologies, with the aim of identifying the cognitive domains that carried the greatest weight in the accelerated aging profile specific to each condition.

## Results

The Ridge regression model trained on the healthy control cohort (N = 394) achieved a correlation of r = 0.942 between chronological age and estimated brain age, with a mean absolute error (MAE) of 3.05 years, in a population ranging from 17 to 64 years of age. In other words, simply by measuring the cognitive performance of a person attending the clinic, the model makes an error of only about three years with respect to chronological age. These values indicate that the 18 digital cognitive metrics from the SPAIn battery accurately capture the normative trajectory of cognitive aging, thereby validating their use as a basis for brain age estimation.

Figure 2 shows a linear relationship between chronological age and estimated brain age across the entire age range of the sample (17–64 years), with homogeneous dispersion suggesting that the model is stable in both younger adults and older individuals within the studied range.

**Figure 1.**
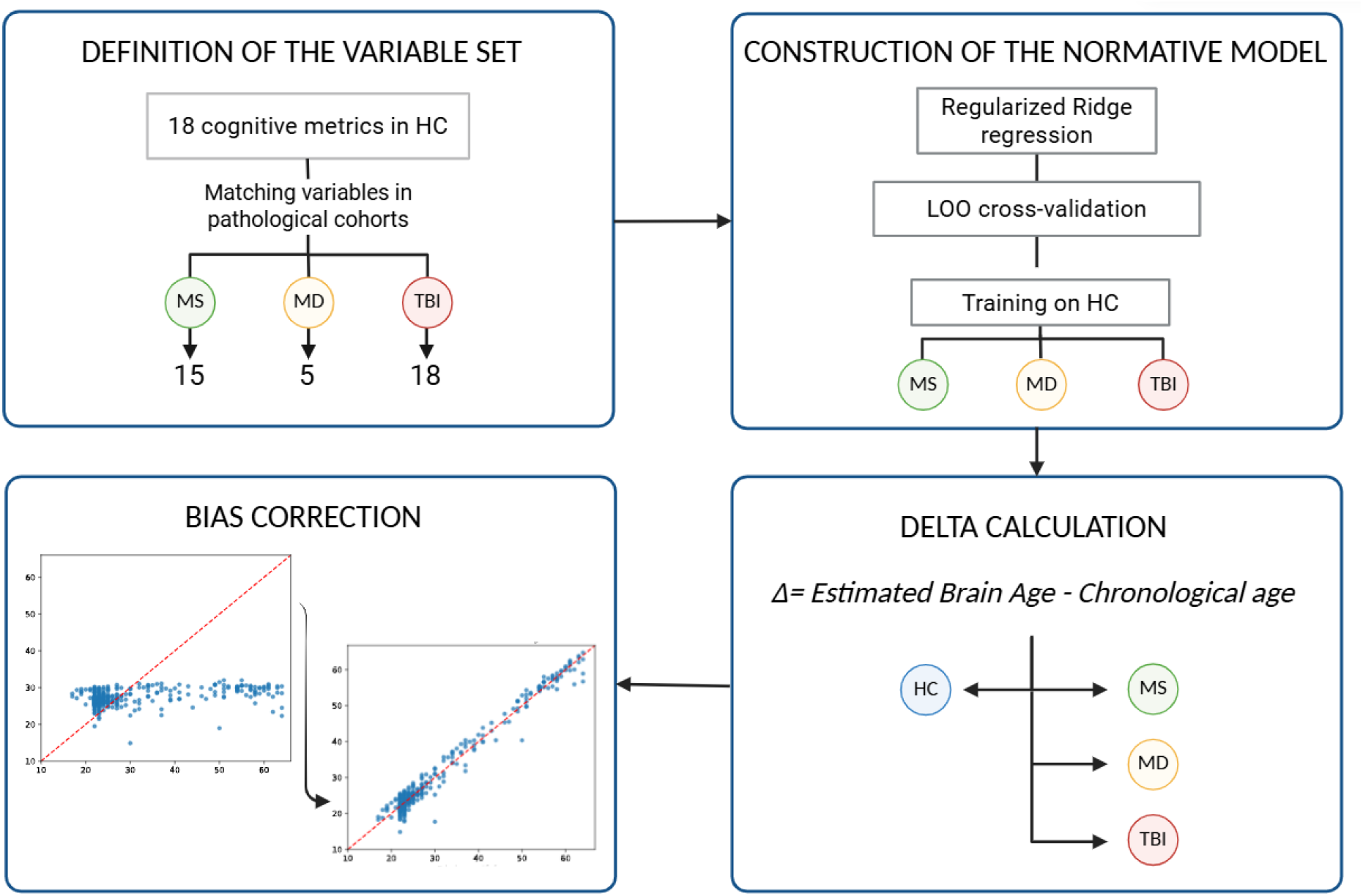
Multicognitive brain age estimation pipeline. (A) Definition of the variable set: the 18 cognitive metrics from the SPAIn battery were used in full in the HC cohort (N = 394); for each clinical cohort, the overlapping subset of variables was used (18 in TBI, 15 in MS, 5 in MD). (B) Construction of the normative model using regularized Ridge regression with Leave-One-Out cross-validation (LOOCV), trained exclusively on HC. (C) Bias correction of delta following the procedure described in the AgeML framework [17]. (D) Calculation of brain age delta (Δ = estimated brain age − chronological age) in each clinical cohort and comparison with healthy controls.

**Figure 2.**
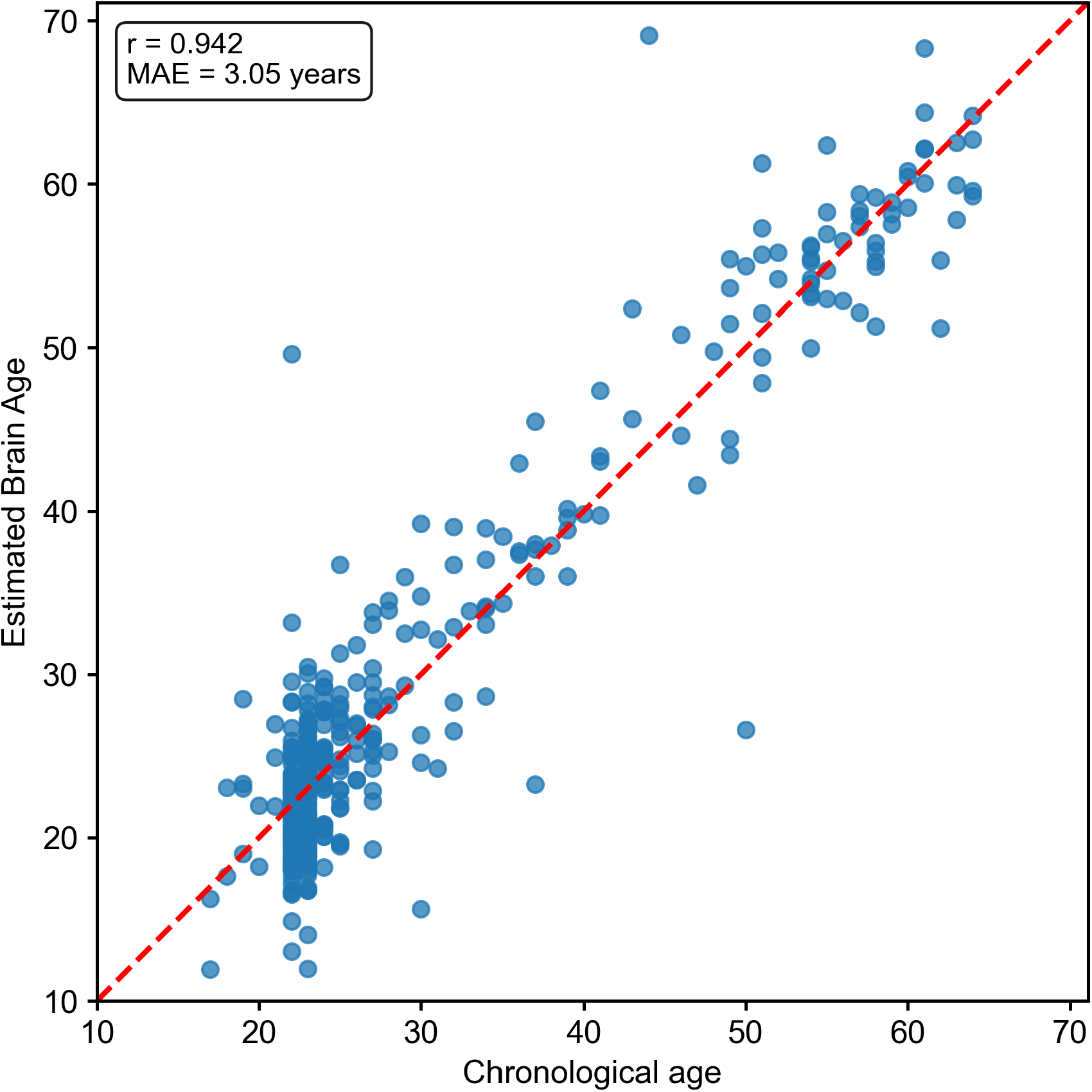
Relationship between chronological age and estimated brain age. (after bias correction) in the healthy control cohort (N = 394). Each point represents one subject. The red dashed line indicates the identity diagonal (brain age = chronological age). The model achieved a correlation of **r = 0.942** with a mean absolute error (MAE) of **3.05 years** in a population aged 17 to 64 years, indicating that the 18 digital cognitive metrics capture the normative trajectory of cognitive aging with high accuracy. The homogeneous dispersion across the entire age range of the sample (17–64 years) suggests model stability in both younger adults and older individuals within the studied age range.

Differences between healthy controls and each clinical group reached statistical significance are shown in Figure 3, confirming the sensitivity of the model to discriminate between normative and pathological aging. Specifically, the traumatic brain injury cohort showed a mean delta of +2.07 years (SD = 7.42; p = 0.004; Cohen’s d = 0.44), the multiple sclerosis cohort showed a mean delta of +1.91 years (SD = 4.68; p < 0.001; d = 0.44), and the depression cohort showed a mean delta of +1.73 years (SD = 1.56; p < 0.001; d = 0.66). In all cases, the differences with respect to healthy controls (Δ ≈ 0 after bias correction) were statistically significant (Mann–Whitney U test, p < 0.01), with small-to-moderate effect sizes according to Cohen’s d, confirming the sensitivity of the model to discriminate between normative and pathological aging.

**Figure 3.**
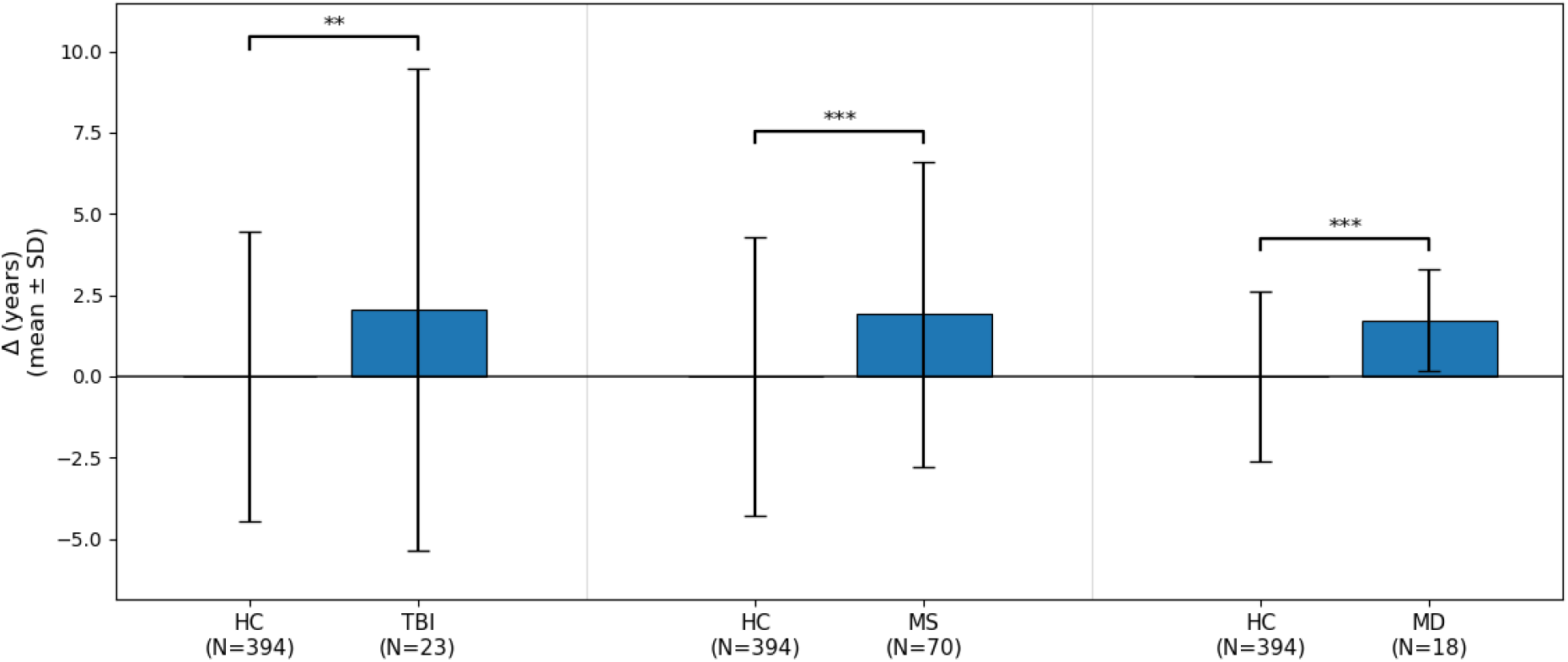
Mean brain age delta (± standard deviation) in healthy controls (HC) versus each clinical cohort: traumatic brain injury (TBI, N = 23), multiple sclerosis (MS, N = 70), and depression (MD, N = 18). Asterisks indicate the level of statistical significance in the Mann–Whitney U test: **p < 0.01; ***p < 0.001. All three clinical cohorts show a significantly elevated delta compared with healthy controls, confirming the presence of accelerated cognitive aging detectable through digital cognitive metrics.

Accelerated cognitive aging was driven in all three clinical cohorts by the same dominant variable: mean reaction time for correct responses in the Simple Reaction Time subtest (PS-02_tmr_successes), reflecting a shared vulnerability in processing speed. However, the degree of concentration of this variable differed markedly across pathologies. In the multiple sclerosis cohort, PS-02_tmr_successes explained 53% of the accelerated aging, pointing to a predominant impairment in elementary processing speed, consistent with previous literature describing slowed information processing as one of the central cognitive alterations in multiple sclerosis [23], [24].

In the traumatic brain injury cohort, PS-02_tmr_successes contributed 27% to delta, representing the single largest contributor but within a notably more distributed profile in which multiple variables from the Go/No-Go and Response Selection subtests also contributed meaningfully. This distributed pattern is consistent with the diffuse nature of traumatic brain injury, which typically affects multiple cognitive systems simultaneously rather than a single domain [25]-[27].

In the depression cohort, PS-02_tmr_successes showed the highest concentration, explaining 79% of delta. This pattern reflects the generalized slowing of information processing—*bradyphrenia*—that constitutes one of the most characteristic cognitive markers of major depression [28], [29].

These results demonstrate that the multicognitive brain age model not only detects accelerated aging, but also reveals the degree to which processing speed concentrates the cognitive deviation in each condition—from a distributed profile in traumatic brain injury to near-exclusive slowing in depression—thereby opening the door to targeted interventions.

## Discussion

The findings of this study suggest that brain age can be estimated with high accuracy from digital cognitive metrics. The Ridge regression model achieved a strong correlation (r = 0.942) between chronological and estimated brain age, with a mean absolute error of only 3.05 years across a sample aged 17–64 years. All three clinical groups showed statistically significant positive deltas relative to healthy controls, and the feature-wise decomposition revealed that processing speed—specifically, simple reaction time—was the dominant contributor to accelerated cognitive aging across all three pathologies, though with markedly different degrees of concentration: 27% in traumatic brain injury, 53% in multiple sclerosis, and 79% in depression. These results extend prior work showing that a neuropsychological brain age model trained on healthy controls could distinguish stable from progressive mild cognitive impairment with high accuracy (AUC = 0.91), and that the resulting delta correlated with time to conversion to Alzheimer’s disease [16]. While that model relied on conventional neuropsychological summary scores (MMSE, ADAS, MoCA, FAQ), the present study employs fine-grained digital metrics—including reaction times and error types—derived from a fully digital battery, which not only estimates brain age but also identifies the specific cognitive domains driving the deviation.

This multidomain approach has direct implications for addressing the challenges of the new longevity. If the goal is not only to live longer, but also to preserve cognition across the lifespan, then tools are needed that can measure, monitor, and eventually improve cognitive health over time [5]. A brain age model based on a digital battery such as SPAIn has several features that make it particularly suitable for this purpose: it is non-invasive, cost-effective, remotely administrable, replicable, and capable of generating immediate and interpretable results.

Unlike neuroimaging-based brain age models, which require costly equipment and hospital infrastructure, a digital cognitive model could be integrated into primary care settings, public health programs, telemedicine platforms, or even digital health applications for home-based monitoring. This scalability is essential if the detection of accelerated cognitive aging is to become not a privilege reserved for those with access to specialized services, but a resource available to the population as a whole.

Beyond detection, the main strength of the multidomain approach proposed here lies in its potential to guide personalized neuropsychological interventions. Although processing speed emerged as the universal dominant contributor to accelerated aging, the degree of concentration varied substantially across conditions: from a distributed profile in traumatic brain injury (27%), where multiple cognitive variables contributed meaningfully, to a near-exclusive concentration in depression (79%), where psychomotor slowing accounts for virtually all of the deviation. This gradient suggests that interventions should be calibrated differently: whereas depression may benefit primarily from targeted processing speed training, traumatic brain injury may require broader cognitive stimulation addressing multiple domains simultaneously.

This finding connects directly with the concept of rejuvenating neuropsychological interventions: cognitive stimulation programs designed not in a generic way, but calibrated according to the concentration profile of each individual. In a patient whose brain age delta is dominated by a single variable (as observed in depression), the intervention would prioritize exercises aimed at rapid information processing and attentional efficiency. In a more distributed profile (as observed in traumatic brain injury), the program would combine processing speed training with executive control and sustained attention tasks.

Digital cognitive rehabilitation platforms—such as NeuronUP, which offers more than 15,000 cognitive stimulation activities organized by domain—provide the technological support necessary to implement this type of personalized intervention at scale. The combination of a brain age model that identifies the most affected domains and an intervention platform that allows for a full assessment-intervention-reassessment loop with strong potential for active aging.

These findings are consistent with the broader shift toward digital innovation in neuropsychological assessment. Recent reviews have highlighted how emerging digital technologies—including ecological momentary assessment, passive smartphone sensors, and wearable devices—can overcome key limitations of traditional assessment, such as low-frequency data collection and limited ecological validity [18]. The brain age model proposed here, built on a remotely administrable digital battery, aligns with this vision and could be further enriched by integrating continuous real-world data streams. Similarly, research within the Boston Process Approach has demonstrated that digital technology can capture kinematic, time-based, and graphomotor parameters that reveal underlying neurocognitive processes not detectable through traditional summary scores [19]. This process-level perspective is particularly relevant to the present work, as the multidomain brain age model identifies domain-specific profiles of accelerated aging—effectively constituting a process analysis at a population scale. The future integration of digital process metrics with predictive brain age models could enhance both early detection and the design of truly personalized cognitive interventions.

Within the framework of active and healthy aging policies promoted by international organizations [30], the integration of cognitive brain age models into population screening programs constitutes a promising avenue. The concept of brain age delta—expressed in an intuitive magnitude, namely how many “extra” or “fewer” years the brain’s functioning appears to have aged—facilitates communication with both the assessed individual and non-specialist professionals, overcoming the interpretative barriers posed by traditional neuropsychological scores.

A screening program based on this model could identify individuals with an elevated positive delta—that is, with faster-than-expected cognitive aging—before they develop subjective cognitive complaints or functional impairment, thereby enabling early preventive interventions. Likewise, periodic administration of the assessment would make it possible to monitor the evolution of delta over time, objectively evaluating the effects of implemented interventions.

Several limitations of the present study should be acknowledged. First, the sample size of the clinical cohorts is small, particularly in traumatic brain injury (N = 23) and depression (N = 18), which limits statistical power and the generalizability of the described impairment profiles. Second, the cross-sectional design does not allow conclusions regarding the temporal evolution of delta or its sensitivity to change following interventions. Third, the absence of a specific cohort of advanced normative aging (individuals older than 65 years without diagnosed pathology) prevents direct validation of the model in the age range of greatest relevance to the new longevity. Finally, generalization to populations with diverse sociocultural and educational backgrounds requires further study.

Future research lines include longitudinal studies assessing the sensitivity of delta to change (for example, after a cognitive stimulation program), extension of the model to cohorts of advanced normative aging, combination of digital cognitive metrics with neuroimaging data in multimodal models, and integration of the model into digital health platforms for use in primary care and population screening contexts.

## Conclusions

Digital cognitive metrics obtained through computerized neuropsychological assessment make it possible to estimate a multidomain brain age with high accuracy (r = 0.942; MAE = 3.05 years). Thus, by assessing only a person’s cognitive performance—including any new individual attending a consultation for screening—the model estimates age with an approximate margin of error of only three years relative to chronological age. This enables robust modeling of both normal and accelerated aging, thereby facilitating the recommendation of optimal rejuvenating strategies for mental health.

To validate our model, we demonstrated accelerated cognitive aging in clinical cohorts of multiple sclerosis, traumatic brain injury, and depression, and showed that processing speed is the universal dominant contributor, with varying degrees of concentration across pathologies. This profiling capacity goes beyond the mere detection of impairment and opens the door to the design of personalized neuropsychological interventions calibrated to the concentration profile of each individual.

In the context of the new longevity, this approach emerges as a promising tool for three complementary objectives: accessible population screening of cognitive aging, the design of individualized rejuvenating interventions, and the objective evaluation of the effectiveness of such interventions over time. Digital multicognitive brain age ultimately represents a step toward a longevity that is not only longer, but also cognitively fuller.

---

**Table 1.** Demographic characteristics of the study cohorts.

| **Demographic characteristics** | **HC** | **MS** | **TBI** | **MD** |
| --- | --- | --- | --- | --- |
| N | 394 | 70 | 23 | 18 |
| Mean age ± SD (years) | 30.0 ± 12.5 | 41.2 ± 8.0 | 32.7 ± 12.7 | 45.5 ± 8.1 |
| Age range (years) | 17–64 | 26–60 | 17–64 | 31–58 |
| Sex (women/men) | 307/87 | 47/23 | 17/6 | 13/5 |
| Predominant educational level | University (68.5%) | — | — | — |

HC: healthy controls; MS: relapsing-remitting multiple sclerosis; TBI: traumatic brain injury; MD: major depression. SD: standard deviation. For the clinical cohorts, educational level was recorded in years of formal schooling (see main text for details).

**Table 2.** Digital cognitive metrics from the SPAIn battery used in the brain age model.

| **Variable** | **Subtest** | **Description** |
| --- | --- | --- |
| PS-01_left_taps | Finger Tapping (left) | Mean number of taps in 10 seconds, left hand |
| PS-01_right_taps | Finger Tapping (right) | Mean number of taps in 10 seconds, right hand |
| PS-01_left_time_reaction | Finger Tapping (left latency) | Mean inter-tap interval, left hand |
| PS-01_right_time_reaction | Finger Tapping (right latency) | Mean inter-tap interval, right hand |
| PS-02_successes | Simple Reaction Time | Number of correct responses |
| PS-02_omissions | Simple Reaction Time | Number of omissions |
| PS-02_tmr_successes | Simple Reaction Time | Mean reaction time for correct responses |
| PS-03_successes | Go/No-Go | Number of correct Go trials |
| PS-03_false_alarms | Go/No-Go | Number of false alarms in No-Go trials |
| PS-03_omissions | Go/No-Go | Number of omissions in Go trials |
| PS-03_tmr_successes | Go/No-Go | Mean reaction time for correct Go responses |
| PS-04_successes | Response Selection | Number of correct responses |
| PS-04_commissions | Response Selection | Number of commission errors |
| PS-04_omissions | Response Selection | Number of omissions |
| PS-04_tmr_successes | Response Selection | Mean reaction time for correct responses |
| PS-04_tmr_commissions | Response Selection | Mean reaction time for commission errors |
| PS-04_tmr_post_error_slowing | Response Selection | Post-error slowing in reaction time |
| PS-04_errors | Response Selection | Total number of errors |

RT: reaction time. The metrics cover performance indicators (correct responses, omissions, commission errors), speed (mean reaction times), and post-error regulation. For the clinical cohorts, the subset of available variables was used: 18 in TBI, 15 in MS, and 5 in MD (see Section 3.2 for details).

**Table 3.**
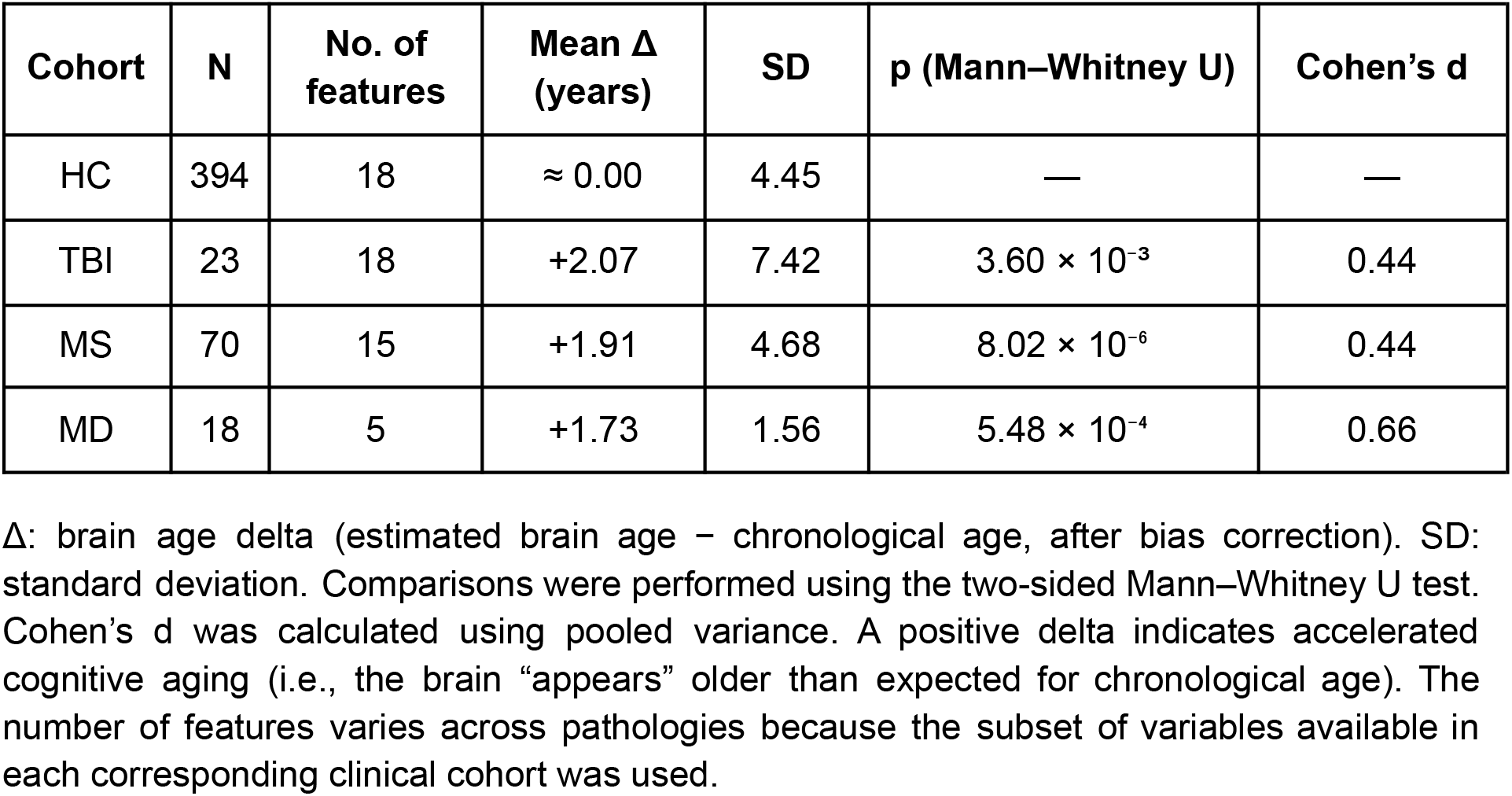
Brain age delta (Δ) by cohort and results of the comparison with healthy controls.

| Cohort | N | No. of features | Mean $\Delta$ (years) | SD | p (Mann–Whitney U) | Cohen’s d |
| --- | --- | --- | --- | --- | --- | --- |
| HC | 394 | 18 | $\approx 0.00$ | 4.45 | — | — |
| TBI | 23 | 18 | +2.07 | 7.42 | $3.60 \times 10^{-3}$ | 0.44 |
| MS | 70 | 15 | +1.91 | 4.68 | $8.02 \times 10^{-6}$ | 0.44 |
| MD | 18 | 5 | +1.73 | 1.56 | $5.48 \times 10^{-4}$ | 0.66 |

$\Delta$ : brain age delta (estimated brain age – chronological age, after bias correction). SD: standard deviation. Comparisons were performed using the two-sided Mann–Whitney U test. Cohen’s d was calculated using pooled variance. A positive delta indicates accelerated cognitive aging (i.e., the brain “appears” older than expected for chronological age). The number of features varies across pathologies because the subset of variables available in each corresponding clinical cohort was used.

**Table 4.** Cognitive variable with the greatest contribution to accelerated aging delta in each pathology.

| **Pathology** | **Dominant variable in $\Delta$** | **Contribution (%)** | **Impaired domain** |
| --- | --- | --- | --- |
| TBI | PS-02_tmr_successes | 27% | Processing speed / slowing |
| MS | PS-02_tmr_successes | 53% | Processing speed / slowing |
| MD | PS-02_tmr_successes | 79% | Processing speed / slowing |

Contribution (%) was calculated by feature-wise decomposition of the differential delta between the clinical cohort and healthy controls, based on the Ridge model coefficients and differences in the standardized means of each variable (see Section 3.4 for details). Impaired domains were assigned according to the SPAIn battery structure. PS-02\_tmr\_successes: mean reaction time for correct responses in the Simple Reaction Time subtest.

